# Retinal optic flow during natural locomotion

**DOI:** 10.1101/2020.07.23.217893

**Authors:** Jonathan Samir Matthis, Karl S Muller, Kathryn Bonnen, Mary M Hayhoe

## Abstract

We examine the structure of visual motion on the retina during natural locomotion in real world environments. Natural locomotion generates a rhythmic translation and rotation profile of the head in space, which means that visually specified heading varies throughout the gait cycle. This presents a challenge if optic flow is to be used to control heading towards a goal. The complex, phasic head movements that occur through the gait cycle create a highly unstable pattern of flow relative to the head. In contrast, vestibular-ocular-reflex mediated fixation simplifies patterns of optic flow on the retinae, resulting in regular features that may be valuable for the control of locomotion. In particular, the sign and magnitude of foveal curl in retinal flow fields specifies the body’s trajectory relative to the fixation point. In addition, the peak in the divergence of the retinal flow field specifies the walker’s instantaneous overground velocity/momentum vector in retinotopic coordinates. Assuming that walkers can determine the body position relative to fixation, this time-varying retinotopic cue for the body’s momentum could provide a visual control signal for foot placement over complex terrain. In contrast, the temporal variation of heading is large enough to be problematic for use in steering towards a goal. Consideration of optic flow in the context of natural locomotion therefore suggests a re-evaluation of the role of optic flow in the visual control of locomotion.

## 1 Introduction

For a mobile creature, accurate vision is predicated on fixation (Land, 2018). Vertebrate photoreceptors are relatively slow, with cones taking up to 20 ms to respond to changes in light intensity (Friedburg, Allen, Mason, & Lamb, 2004). As such, sighted animals must stabilize their eyes relative to the external world to resolve an image that may be used for the control of action. This may be the reason that nearly all vertebrates utilize a “saccade and fixate” strategy whereby gaze is rapidly moved to a new location and then kept stable by way of a gaze stabilization reflexes such as the vestibular ocular reflex (VOR) (Dietrich & Wuehr, 2019). These gaze stabilization mechanisms are phylogenically ancient, with evidence for compensatory eye movements stretching back to the origin of bony fish approximately 450 million years ago (Land, 2018; Walls, 1962). These reflexes are extremely low-level, such that (excepting saccades) it is not possible for a healthy human to move their body without stabilizing their retinae relative to some point in the external world. This oculomotor strategy, along with the highly foveated nature of the primate vision, means that the human visual system is fundamentally built on our ability to rapidly direct gaze to points of interest in the world and then fixate that point as we move our bodies through space.

This movement of the of the body through space creates image motion on the retinae. In the absence of eye movements, an observer moving through a structured environment will experience a pattern of radial expansion centered on the Focus of Expansion (FoE), that specifies the current heading direction (Gibson, 1950, 1979). Subsequent work showed that observers are able to judge their heading in simulated flow fields with very high accuracy, on the order of 1-2 degrees of error (W. H. Warren, 1988). As a consequence of this observation, together with a large body of related work, it became generally accepted that the optic flow patterns arising from linear translation are used by observers to control their direction of heading during locomotion (see reviews by Lappe, Bremmer, & van den Berg, 1999; W. H. Warren, 2007; Li & Cheng, 2013; Britten, 2008). Similarly, this idea has dominated the interpretation of neural activity in motion-sensitive areas of visual cortex such as MST and VIP, areas that respond selectively to optic flow stimuli and are thought to underlie both human and monkey heading judgments (Britten, 2008; A. Chen, Gu, Liu, DeAngelis, & Angelaki, 2016; X. Chen, DeAngelis, & Angelaki, 2013; Duffy & Wurtz, 1991; Morrone et al., 2000; Wall & Smith, 2008; Bremmer, Churan, & Lappe, 2017; Kaminiarz, Schlack, Hoffmann, Lappe, & Bremmer, 2014).

One complication of this simple strategy is that whenever an individual fixates anywhere but the instantaneous direction of travel, the singularity of the retinal flow field is located at the gaze point rather than the direction of heading. By definition, the act of fixation nulls image at the fovea, so retinal flow during fixation on a stable point in a static scene will always involve patterns of outflowing motion from null motion at the fovea. The resulting flow field will comprise both a rotational and a translational component, and the question of how walkers might recover their heading direction from these retinal optic flowfields has been been central to much of the research on optic flow (dubbed the “rotation problem” (e.g. Britten, 2008), or the “eye movement problem” (W. H. Warren & Hannon, 1990)). In the large body of subsequent research, it was definitively shown that subjects can make accurate judgements their direction of heading during eccentric fixation, in a process believed to involve using a combination of oculomotor and retinal information to recover the translational (heading) component of retinal optic flow (Longuet-Higgins & Prazdny, 1980; Heeger & Jepson, 1992; Perrone, 1992; Royden, 1997; Wang & Cutting, 1999; Perrone, 2018)

However, not matter what the eyes do during locomotion, the complex, phasic movements of the head throughout the bipedal gait cycle will strongly increase the variability in the optically specified instantaneous heading direction. The stimuli typically used in studies of optic flow simulate constant velocity straight-line movement. These stimuli produce a strong sense of self-motion, or vection, and are a reasonable approximation to the airplanes, automobiles, and gliding birds that were the basis of Gibson’s original insights. However, natural locomotion results in rhythmic translation and rotation profiles of the head in space, peaking at approximately 2Hz, which is the natural period of footsteps (Menz, Lord, & Fitzpatrick, 2003; Kavanagh & Menz, 2008). This means that the momentary heading direction varies through the gait cycle, creating a complex pattern of flow on the retina. While this has long been recognized (e.g. Lappe & Hoffmann, 2000; Calow & Lappe, 2007, 2008),the gait induced oscillations are typically assumed to be a minor factor and are referred to as “jitter” (e.g. van den Berg & Beintema, 2000; Nakamura, Palmisano, & Kim, 2016; Palmisano, Gillam, & Blackburn, 2000). During natural locomotion, gaze patterns vary depending on the complexity of the terrain (Matthis, Yates, & Hayhoe, 2018). The gait-induced instabilities of the head and the terrain-dependent patterns of eye movements during natural locomotion determine the actual flow patterns on the retina. There has been some exploration of retinal motion patterns by measuring eye and head movements during locomotion (Einhöuser et al., 2007; Calow & Lappe, 2008). However, those measurements focused on the statistics of retinal flow, rather than the time-varying evolution of the signals. Many algorithms exist to recover observer translation and rotation from the instantaneous retinal flow field. However, there is disagreement on whether heading perception is based on the instantaneous flow field, or instead based on some measure of the way the flow patterns change over time (Cutting & Springer, 1992; W. H. Warren, Blackwell, Kurtz, Hatsopoulos, & Kalish, 1991; Burlingham & Heeger, 2020; Li & Cheng, 2011).

The goal of this paper is to measure eye, body, and head movements during natural locomotion and to use this data to investigate the resulting optic flow patterns. We first calculated the flow patterns relative to the head, as this reflects the way that the movement of the body during gait impacts instantaneous heading direction by showing an eye-movement-free representation of optic flow. Then, we combine these head-centered flowfields with measured eye position to estimate the retinal optic flow experienced during natural locomotion. By characterizing the optic flow stimulus experienced during natural locomotion, we may gain a greater insight into the ways that the nervous system could exploit these signals for locomotor control.

We recorded subjects walking in natural environments while wearing a full-body, IMU-based motion capture suit (Motion Shadow, Seattle, WA, USA) and a binocular eye tracker (Pupil Labs, Berlin, Germany). Data from the two systems were synchronized and spatially calibrated using methods similar to those described in Matthis et al. (2018) (See Methods for details). This generated a data set consisting of spatiotemporally aligned full-body kinematic and binocular gaze data (sampled at 120Hz per eye), as well as synchronized first-person video from the head-mounted camera of the eye tracker (100 deg. diagonal, 1080p resolution, 30Hz). The integrated data are shown by the skeleton and gaze vectors in Figure 1B and Video 1. Subjects walked in two different environments: a flat tree-lined trail consisting of mulched woodchips shown in Figure 1 (selected because it was highly visually textured, but flat enough that foot placement did not require visual guidance), and a rocky, dried creek bed, where successful locomotion requires precise foot placement (This was the same terrain used in the ‘Rough’ condition of Matthis et al. (2018)). Subjects walked the woodchips path while completing one of three experimental tasks: “Free Viewing,” where subjects were asked to walk while observing their environment with no explicit instruction; “Ground Looking” where subjects were asked to look at the ground at a self-selected distance ahead of them (intended to mimic gaze behavior on the rocky terrain without the complex structure or foothold constraints)^1^.; “Distant Fixation”, wherein subjects were asked to maintain fixation on a self-selected distant target that was roughly at eye height (this condition was intended to most closely mimic psychophysical tasks that have often been employed to explore perception of heading, (e.g. Lappe et al., 1999; Paolini, Distler, Bremmer, Lappe, & Hoffmann, 2000; Royden, Banks, & Crowell, 1992; W. H. Warren & Hannon, 1990). Walking over the rocky terrain (labelled “Rocks”) was demanding enough that subjects were not given instructions other than to walk from the start to the end position at a comfortable pace.

**Figure 1.**
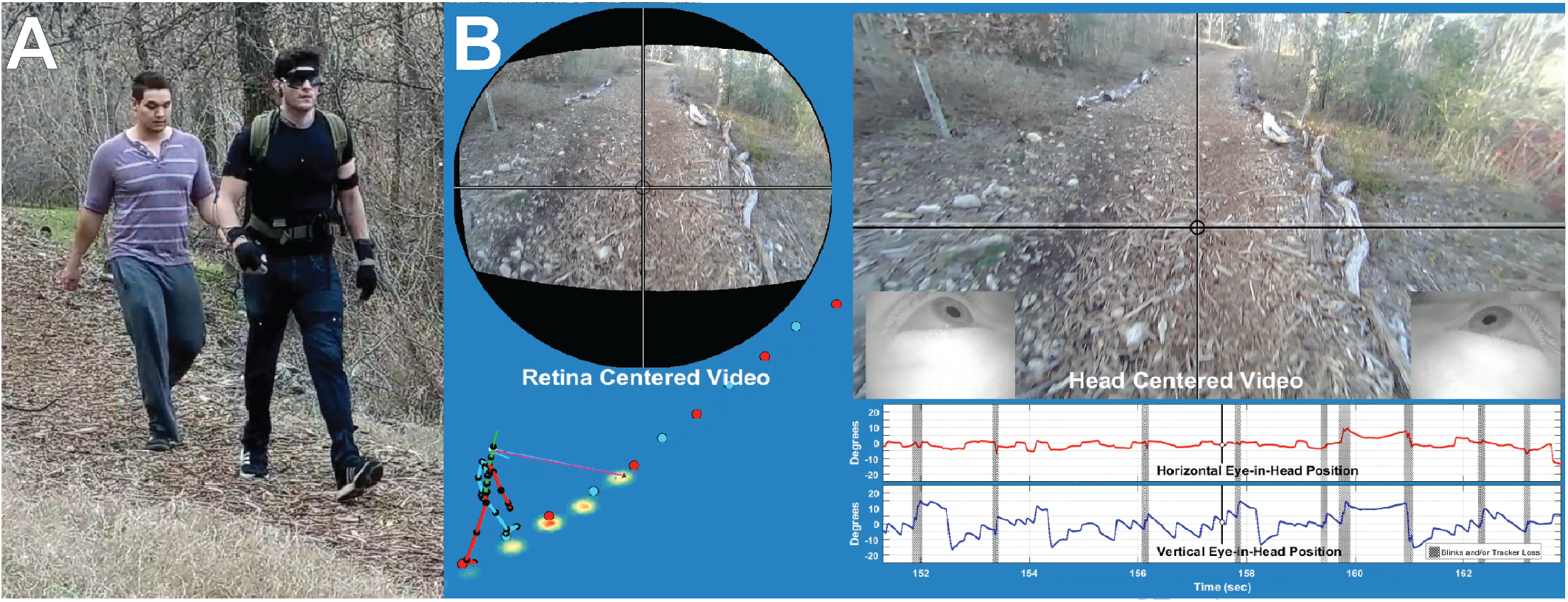
The data collection setup. (A) shows the subject walking in the Woodchips terrain wearing the Pupil Labs binocular eye tracker and Motion Shadow motion capture system. Optometrist roll-up sunglasses were used to shade the eyes to improve eye tracker performance. (B) shows a sample of the data record, presented as a sample frame for Video 1. On the right is the view of the scene from the head camera, with gaze location indicated by the crosshair. Below that are the horizontal and vertical eye-in-head records, with blinks/tracker losses denoted by vertical gray bars. The high velocity regions (steep upwards slope) show the saccades to the next fixation point, and the lower velocity segments (shallow downwards slope) show the vestibular ocular reflex component that stabilizes gaze on a particular location in the scene as the subject moves towards it, resulting a characteristic saw-tooth appearance for the eye signal (without self-motion and VOR these saccades would exhibit a more square-wave like structure). On the left, the stick figure shows the skeleton figure reconstructed form the Motion Shadow data. This is integrated with the eye signal which is shown by the blue and pink lines. The representation of binocular gaze here shows the gaze vector from each eye converging on a single point (the mean of the two eyes). The original ground intersection of the right and left eye is shown as a magenta or cyan dot (respectively, more easily visible in Video 1). The blue and red dots show the foot plants recorded by the motion capture system. The top left figure shows the scene image centered on the point of gaze reconstructed from the head camera as described in the Methods section.

## 2 Results

### 2.1 Optic flow during real-world locomotion

#### 2.1.1 Effect of gait on head-centered optic flow

To measure the optic flow patterns induced by the body motion independently of eye position, we first ran the videos from the head-mounted camera of the eye trackers through a computational optic flow estimation algorithm (DeepFlow, Weinzaepfel, Revaud, Harchaoui, & Schmid, 2013), which provides an estimate of image motion for¿ every pixel of each frame of the video (Video 2). As an index of heading we tracked the focus of expansion (FoE) within the resulting flow fields using a novel method inspired by computational fluid dynamics (See Methods). This analysis provides an estimate of the FoE location in head-centered coordinates for each video frame (Video 3).

We found that the head-centered optic flow pattern is highly unstable, rarely corresponds to the walker’s direction of travel, and never lies in a stable location for more than a few frames at a time. The FoE moves rapidly across the camera’s field of view at very high velocities, with a modal velocity across conditions of about 255 deg per sec. Note that this analysis is entirely based on the head-mounted camera and does not rely on the IMU measurements. To ensure that this instability is not just a consequence of the video-based technique, we also measured the FoE in the video of a gimbal-stablized quadcopter camera drone (DJI Phantom 4, Nanshan, Shenzhen, China) and found it to be stable (See Methods and Video 14). Thus the motion of the FoE is not an artifact of the method used here.

The instability of the head-centered flow can be seen in Figure 2, which shows the head-centered optic flow over a period of 150ms (5 frames at 30fps). The red line traces the movement of the FoE over the 5 frames (150ms). The inset in the figure shows the velocity distribution for the FoE, where the average of the 4 conditions is shown in black, and the distributions for the different conditions is shown by the colored curves. Note that the instability of the FoE is relatively unaffected by either the terrain (rocks or woodchips) or by subject’s choice of gaze point, which is linked with head posture. The instability of the head-centered FoE is also clearly demonstrated in Video 3, which shows the integral lines of the optic flow fields shown in Video 2 with the FoE on each frame denoted by a yellow star. This lack of stability means that heading during locomotion is more complex than simply accounting for eye position and such a mechanism is only viable for constant velocity motion. Head trajectory during locomotion cannot be approximated by constant velocity, straight line motion or with added ‘jitter’ (Palmisano et al., 2000).

**Figure 2.**
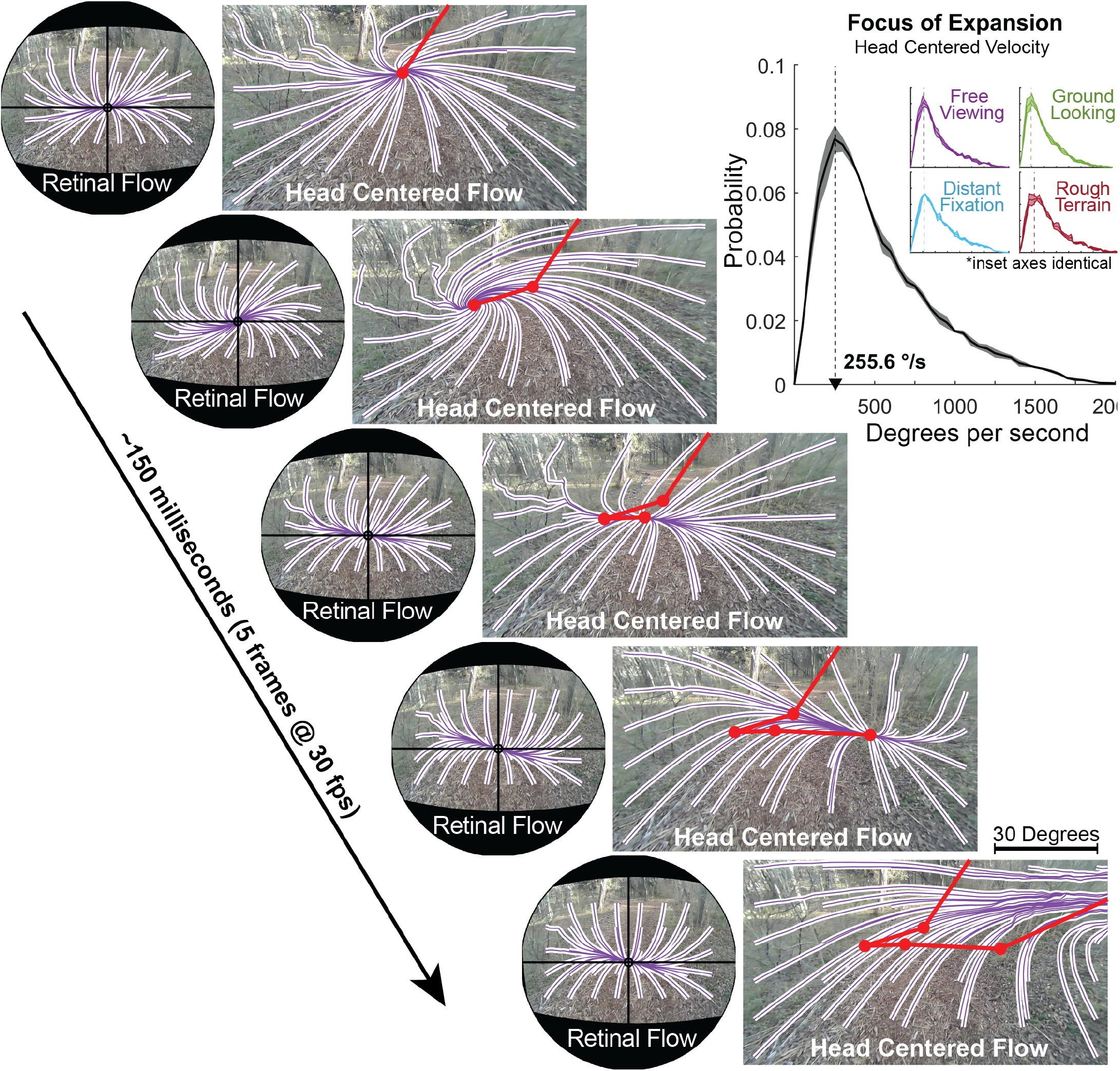
Retinal vs Head-Centered Optic flow. Optic flow patterns (down-sampled) for a sequence of 5 video frames from Video 3, altered for visibility in print. Head centered flow shows optic flow in the reference frame of the head mounted “world” camera, and represents optic flow free from the effects of eye movements. Retinal flow shows optic flow in the references frame of a spherical pinhole camera stabilized on the subject’s fixation point. Purple and white lines show the integral curves of the measured flow fields, determined by using the streamlines2 function in Matlab to make a grid of particles drift along the negation of the flow fields measured by the DeepFlow optical flow detection algorithm in OpenCV. The red trace shows the movement of the head-centered FoE moving down and to the right across the 5 frames. Note the stability of the retina-centered flow in contrast. The inset shows the probability of FOE velocity across all conditions (black histogram), as well as split by condition (colored insets). The thick line shows the mean across subjects, and shaded regions show +/−1 standard error.

#### 2.1.2 Head velocity oscillations during natural locomotion

The instability of head-centered optic flow arises from the natural oscillations of the head during locomotion. A walker’s head undergoes a complex, repeating pattern of velocity throughout the gait cycle, as shown in Figure 3. Although the vestibulo-collic and spinocollic reflexes result in some damping of head acceleration in the Anterior/Posterior and Medial/Lateral directions (note the flattening of the acceleration profile between the hips, chest, and head in these dimensions), no such damping appears to occur in the vertical direction, most likely because it would interfere with the efficient exchange of potential and kinetic energy inherent in the inverted pendulum dynamics of the bipedal gait cycle (Donelan, Kram, & Kuo, 2001; Kuo, 2007; Mochon & McMahon, 1980). As a result, a walker’s head undergoes a continuously changing phasic velocity profile over the course of each step (see yellow vector and trace in Video 3, which shows the velocity vector of the walker’s eyeball center at each frame of the recording). The peak magnitude of these velocity oscillations (particularly in the vertical dimension) is approximately half the walker’s overall walking speed, as shown in the Figure.

**Figure 3.**
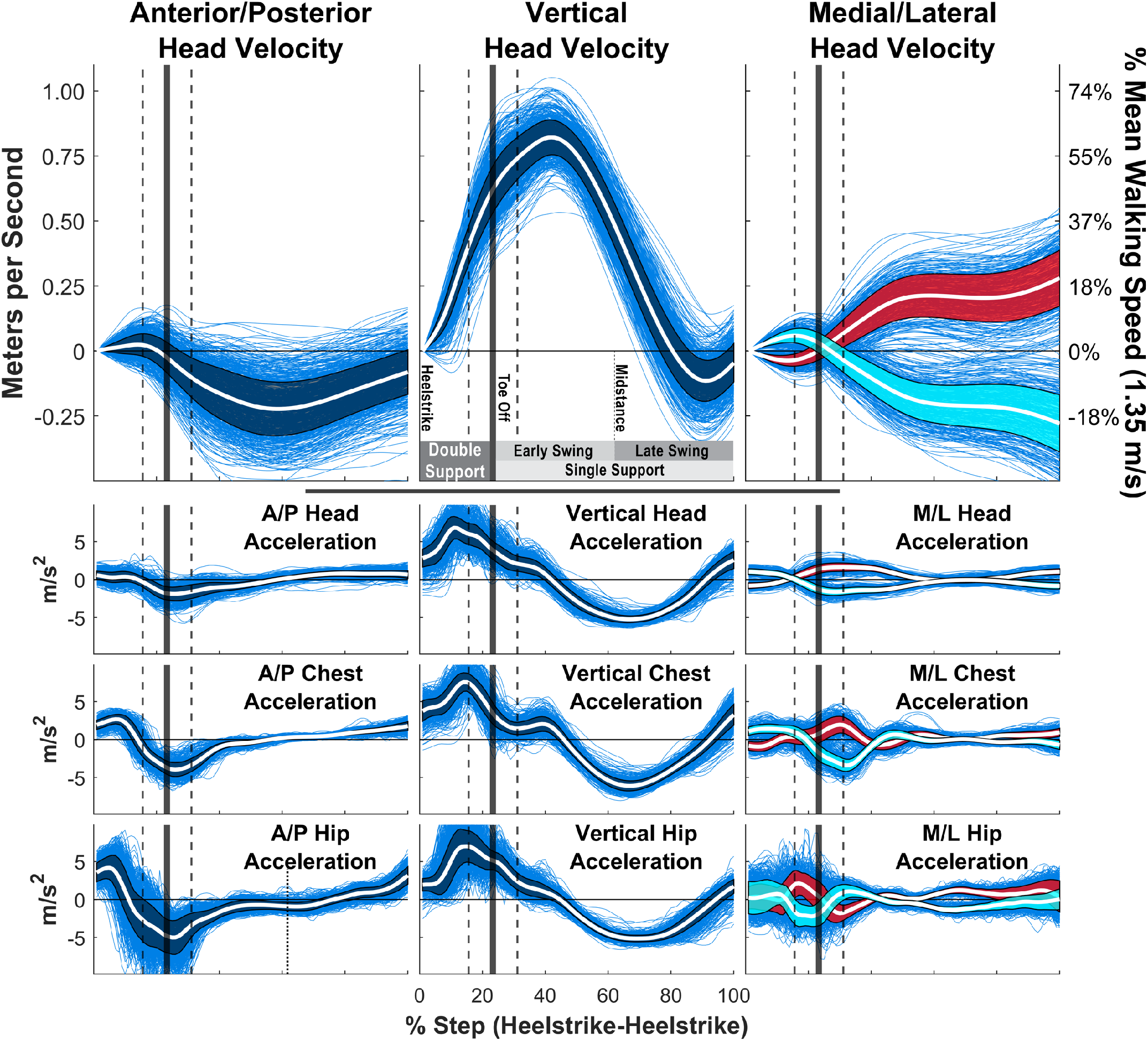
Head Velocity and Acceleration during locomotion. Head velocity and head/chest/hips acceleration patterns of a single representative subject walking in the ‘‘Distant Fixation” condition (where they walk while maintaining fixation on a point far down their path). Each blue trace is the velocity or acceleration trace for a single step, normalized for the time period from heel-strike to the subsequent heel-strike. The white line shows the mean and the shaded region is +/−1 standard deviation. Right and Left steps were computed separately for the Medial/Lateral data, and are shown in red and cyan respectively. The vertical solid lines show mean toe-off time, with the vertical dashed lines showing +/−1 standard deviation. Other subjects and conditions can be found in the Supplemental Information.

These variations in head velocity (Figure 3) and the accompanying rotations (Supplemental Figure 4) throughout the gait cycle are the reason why the head-centered flow patterns move at such high velocities across a walker’s visual field. The location of the FoE at any given moment will be defined by the head’s velocity vector (Longuet-Higgins & Prazdny, 1980), so changes in that vector will lead to changes in the FoE’s location in the visual field. Furthermore, because the FoE arises from the visual motion of objects in the walker’s field of view, the linear velocity of the FoE in the viewing plane of the walker will be determined by the angular variations in the walker’s head velocity vector projected onto the visible objects within walker’s environment. Consequently, small angular changes in the walker’s head velocity result in rapid, large scale movements in the FoE on the walker’s image plane. While the importance of the FoE itself has been questioned (W. H. Warren & Hannon, 1990; Lappe et al., 1999), the instability applies to the entire flow field.

#### 2.1.3 Retinal optic flow during natural locomotion

To estimate retinal flow patterns during natual locomotion, we first identified fixations in the eye tracker data and then aligned each frame from head-mounted video so that the point of fixation was always pinned to the center of the image (see Methods). Then, the resulting ‘gaze stabilized’ videos were analyzed with the same optic flow and streamline based analysis used for the head-centered videos. A comparison of retinal and head centered flow patterns during a single fixation is shown in Figure 2 It can be seen that the retinal patterns change only modestly over time as a consequence of the gaze stabilization. This suggests that gaze stabilization/fixation may be an important component in understanding the use of optic flow patterns, as has been suggested by Angelaki and Hess 2005, Glennerster et al 2001, and Calow Lappe 2008, rather than a complicating factor as has often been assumed (e.g. Britten, 2008). When gaze is properly stabilized on a location in the world during locomotion, the result will always be a pattern of radial expansion combined with rotation, centered on the point of fixation, this suggests that the critical information is in the structure of the retinal flow pattern. In what follows, we explore some of the task relevant features of retinal flow.

### 2.2 Simulated retinal flow

It is useful to examine the visual motion incident on the walker’s retina, since this is the signal that is directly available to the visual system. To investigate the structure of the retinal motion patterns, or retinal flow, we used the walkers’ recorded gaze and kinematic data to geometrically simulate the flow patterns experienced during natural locomotion.

These simulations of retinal flow utilize a spherical pinhole camera model of the eye (Figure 4). The spherical eye maintains ‘fixation’ on a point on the ground, while a grid of other ground points is projected through the pupil and onto the back of the spherical eye. The locations of these projected ground points on the retina are tracked frame by frame to calculate simulated retinal optic flow. The location of this eyeball can be set to maintain fixation while following an arbitrary trajectory through space to determine the structure of retinal optic flow during various types of movement (Videos 4-9). In addition, this model of the eye can be set to follow the trajectory of walker’s eyes as they walked in natural terrain (Videos 10 and 13) in order to estimate retinal motion experienced during natural locomotion, without the complicating elements associated with the real-world video recordings (see Methods for details).

**Figure 4.**
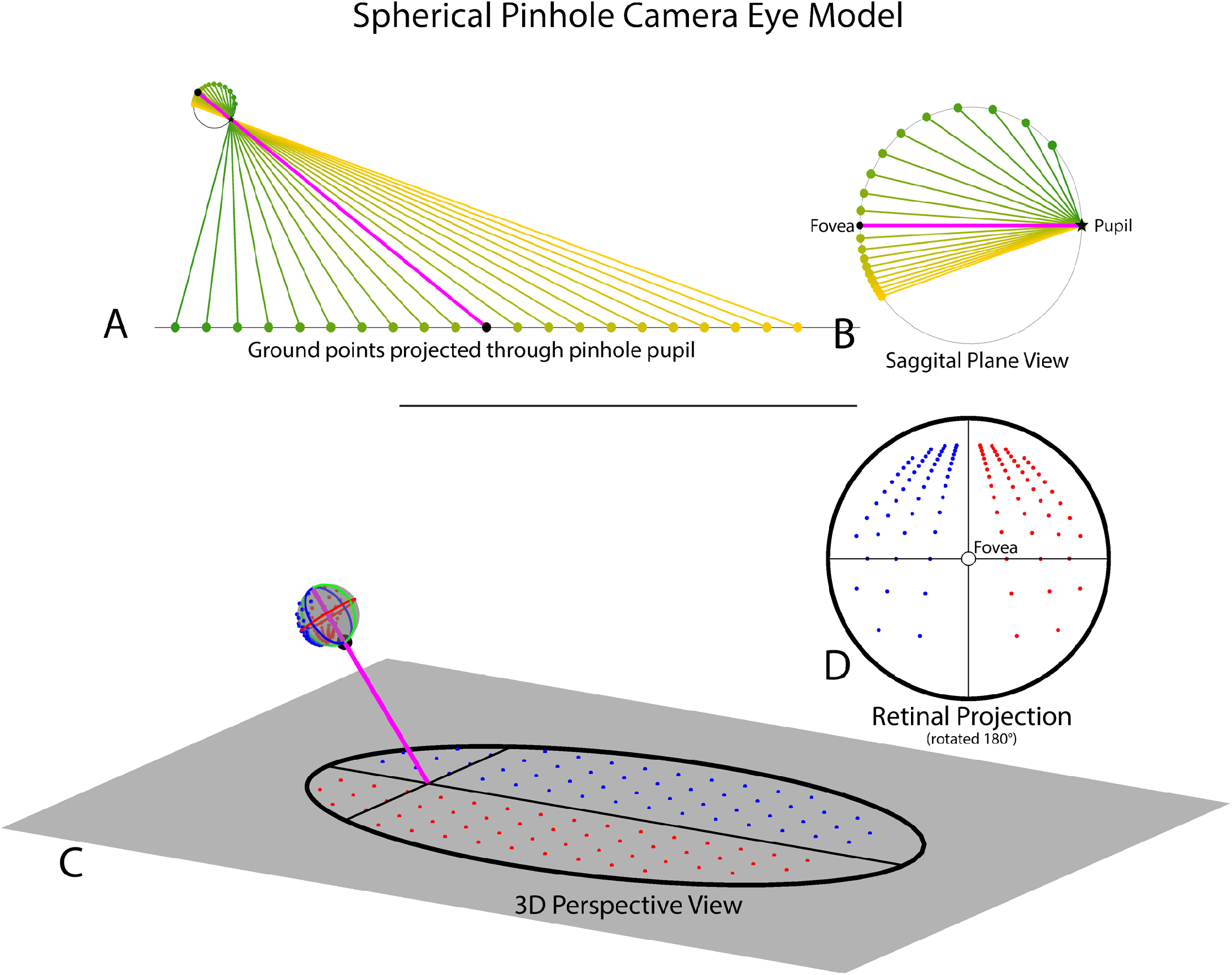
Spherical Pinhole Camera Model of the Eye. The spherical pinhole camera model of the eye used to estimate retinal optic flow experienced during natural locomotion. A, B show a sagittal plane slice of the 3D eye model. A shows the eye fixating on a point on the ground (pink line shows gaze vector, black circle shows fixation point) as points in the upper (orange) and lower (green) visual fields project on the back of the eye after passing through a pinhole pupil. B shows a closer view of the sagittal slice of the eye model. C, D show the full 3D spherical pinhole eye model. C shows the 3D eye fixating a point on the ground (black crosshairs), with field of view (60 degree radius) represented by the black outline. Note that the circular field of view of the eye is elongated due to its projection onto the ground plane. Red and blue dots represent points in the right and left visual field, respectively. D shows the retinal projection of the ground points from C on the spherical eye model. Ground dot location in retinal projection is defined in polar coordinates (ν, ρ) relative to the fovea at (0,0), with ν defined by the angle between that dot’s position on the spherical eye and the ‘equator’ on the transverse plane of the eye (blue circle) and ρ defined as the great-circle (orthodromic) distance between that dot and the fovea of the spherical eye. The retinal projection has been rotated by 180 degrees so that the upper visual field is at the top of the image.

#### 2.2.1 Retinal cues for the control of locomotion

Several features of the retinal flow patterns provide powerful cues for the visual control of locomotion. Stabilization of gaze during fixation nulls visual motion at the fovea, so the basic structure of retinal optic flow will always consist of outflowing motion centered on the point of fixation. The retinal motion results from the translation and rotation of the eye in space, carried by the body and the walker holds gaze on a point on the ground during forward movement.^2^

#### 2.2.2 Foveal curl provides a cue for locomotor heading relative to the fixation point

When a walker fixates a point that is aligned with their current velocity vector, the resulting retinal flow field has no net rotation at the fovea (Figure 5A, Video 4). However, fixation of a point that is off of their current trajectory results in a rotating pattern of flow around the fovea that may be quantified by calculating the curl of the retinal vector field. Fixation of a point that is to the left of the current path results in counter-clockwise rotation (Figure 5B, Video 5), while fixation to the right of the current trajectory results in clockwise rotation around the fovea (Figure 5C, Video 6). This shows that the curl of retinal flow around the fovea provides a metric for the walker’s trajectory relative to their current point of fixation.

**Figure 5.**
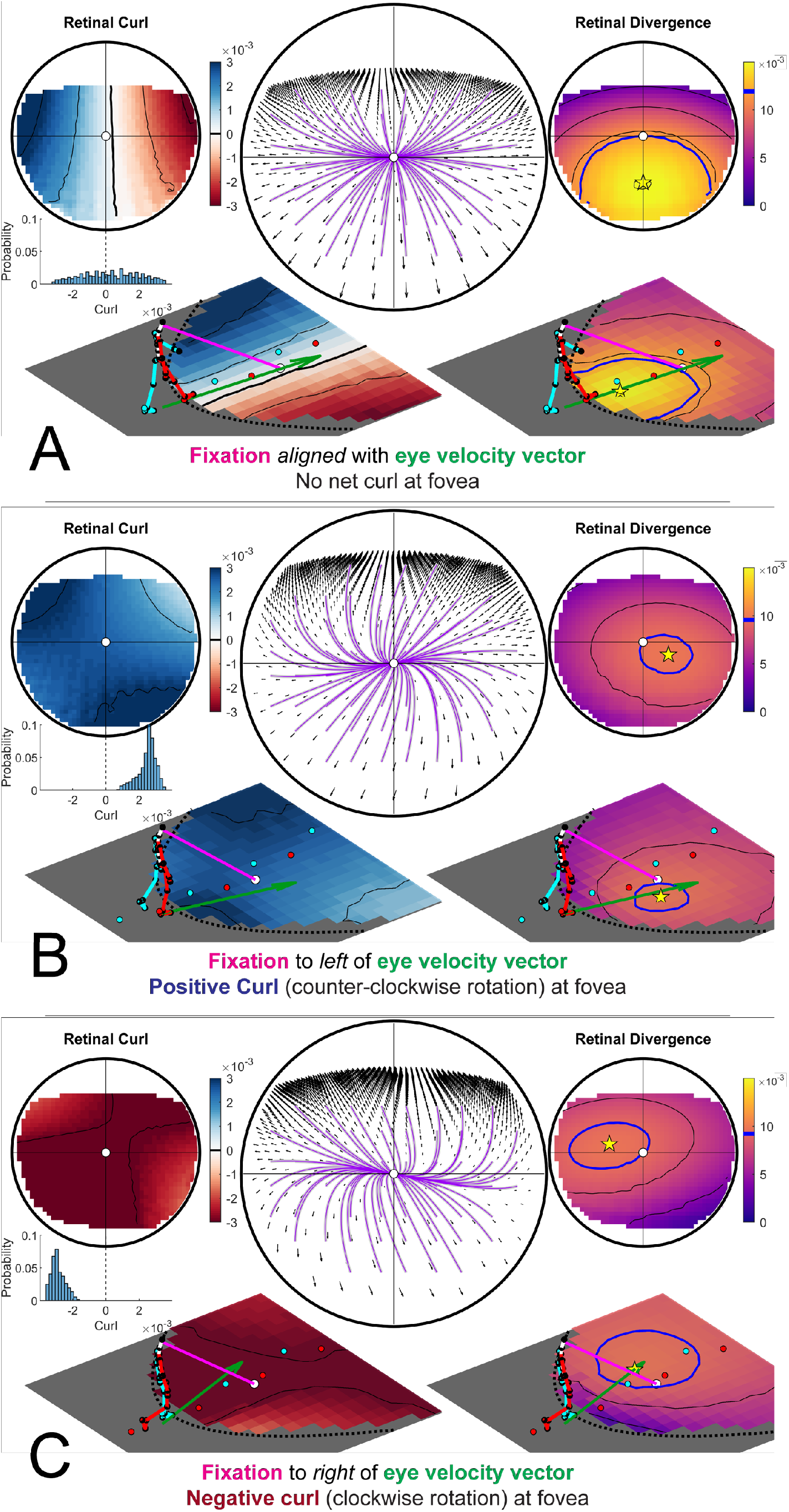
Retinal optic flow during natural locomtoion. Retinal flow simulation based on integrated eye and full-body motion of an individual walking over real world terrain. Panels A-C are based on frames taken from Video2, which shows data derived from a subject walking in the Woodchips condition under the Ground Looking instructions. A shows a case where the fixation point (pink line) is aligned with ground projection of the eye’s velocity vector (green arrow). Middle circular panel shows simulated optic flow based on fixation location and body movement. Left and right circular panels show the results of applying the curl and divergence operators to the retinal flow field in the middle panel. Left and right bottom panels show the projection of the curl (left) and divergence (right) onto a flat ground plane. The green arrow shows the walker’s instantaneous velocity vector (scaled for visibility), which always passes through the maximum of the retinal divergence field, which always lies within the foveal isoline (blue circle). B and C show cases where the fixation point is to the left or right of the eye’s velocity vector (respectively). Fixation to the left of the eye’s velocity vector (B) results in a global counter-clockwise rotation of the retinal flow field and positive retinal curl at the fovea, while fixation to the right of the eye’s velocity vector results in clockwise flow and negative curl at the fovea.

Thus, the magnitude of rotation around the point of fixation provides a quantitative cue for the walker’s current trajectory relative to the point of fixation. Specifically, there will be zero net curl around the fovea when the walker is fixating a point that is aligned with their current trajectory, and positive/negative curl values for fixation to the left/right of the current trajectory, with the magnitude of the curl signal increasing proportional to the eccentricity of the fixation point (Video 8). This cue could be used to help the walker move the body in a particular direction relative to the current fixation location on the ground. Maintaining fixation on a specific location and turning until there is zero curl around the fovea would be an effective way for a walker to orient the body towards a specific location (such as a desirable foothold in the upcoming terrain), correct for potential postural errors, or simply to maintain a straight heading by balancing the horizontal trajectory changes that occur during steps on the right or left foot (Video 10). Similarly, walkers might learn to predict a certain pattern of retinal flow for a given bodily movement, and values outside this range could provide a signal for balance control (Reimann, Fettrow, Thompson, & Jeka, 2018; Roth et al., 2016)

#### 2.2.3 The point of maximum divergence encodes the body’s over-ground momentum vector in retinotopic coordinates

Fixating a point on a point on the ground while moving forward creates a pattern of expansion centered around the point of fixation. The local expansion at each point of a vector field can be quantified by calculating the *divergence* of the vector field. Divergence can be intuitively thought of as the rate that an array of drifitng particles would vacate each point in the flow field. In contrast to optic flow on a fronto-parallel plane, the distance gradient resulting from fixation on the groundplane results in variation in motion parallax of points on the ground plane across the upper to lower visual field. Therefore, divergence of the resulting flow field is determined by the velocity of the eye relative to the fixated point combined with the parallax gradient of points on the ground plane (Koenderink & van Doorn, 1976). In the context of locomotion, the divergence field during a ground plane fixation results in a hill-like gradient with a peak denoting the point of maximum retinal divergence that is contained within the ellipsoidal isoline that passes through the point of fixation (green circle in Figure 5). During straight-line movement towards the point of fixation the foveal iso-ellipsoid begins in the lower visual field before constricting to a point when the observer is one eye height away from their point of fixation (that is, when their gaze angle is 45 degrees) and then expanding into the upper visual field (Figure 5A-C, Video 4-6, Note that this feature of the divergence field is affected by upward and downward trajectories of the eye (Video 7).

Projecting the retinal divergence map onto the ground plane reveals that the ground projection of the eye’s velocity vector always passes through the point of maximum retinal divergence. This feature of retinal flow means that a walker could, in principle, discern their own world-centered ground velocity directly from retinal flow. Thus it should be possible to determine the body’s world-centered velocity vector directly from patterns of motion on the retina.Because the walker’s mass is constant, this velocity vector may serve as a proxy for the body’s momentum vector in retinotopic coordinates. The body’s momentum vector is the information a walker needs to make biomechanically informed decisions about where to place their feet so the ability to derive this information directly from retinal optic flow may be of critical importance for the control of locomotion. Since the walker can determine the body-relative location of their fixation point (Crawford, Henriques, & Medendorp, 2011), this analysis shows how patterns of stimulation on the retina might be directly related to the physical dynamics of bipedal locomotion (Matthis & Fajen, 2013, 2014; Matthis, Barton, & Fajen, 2015, 2017; Barton, Matthis, & Fajen, 2017).

The correspondence of the projection of the eye’s velocity vector onto the ground plane with the point of maximum divergence was previously described by Koenderink and van Doorn, (Koenderink & van Doorn, 1976). The use of this information as a cue for heading was subsequently challenged by Warren and Hannon (W. H. Warren & Hannon, 1990). In one of the behavioral experiments described in that paper, Warren and Hannon asked stationary subjects viewing a computer monitor to judge their heading during simulated motion through a 3D star field. The authors argue that because these star fields were spatially discontinuous, the resulting flow fields will not contain a point of maximum divergence. They took the finding that subjects made accurate heading estimates in these star fields as evidence that humans do not use the point of maximum divergence to determine their heading direction. There are at least two problems with this line of argument. First and foremost, the finding that subjects can make accurate heading judgements in the absence of a particular cue is not evidence that they do not use that cue when it *is* present. Secondly, even if it is the case the the stimuli presented to subjects were sparse or discontinuous to the point where divergence was undefined, this does not mean that the stimuli encoded by subjects would have the same properties. The observer judgments have been shown to be robust to sparse, discontinuous motion stimuli (e.g. P. A. Warren & Rushton, 2009; Rushton & Warren, 2005), so the mathematical properties of the experimental stimuli may not match the stimuli that make it through the complex neurophysiological filters of the visual system. Finally, and relating to the first point, the stimuli used in this study were decidedly unrealistic and do not represent a natural occurring pattern of visual motion, so this study should not be taken as strong evidence against the use of retinal divergence for the control of action. However, it is worth reiterating that the present, purely observational investigation also does not provide any evidence that walkers use divergence in the control of locomotion. Rather, we simply show that the the point of maximum divergence is a reliable feature of fixation-mediated retinal optic flow that appears to encode behaviorally relevant information that *could* be valuable for the control of locomotion. The question of whether walkers actually use this cue - or even if the visual system is physiologically capable of detecting it - must be a matter of experimental investigation.

In addition, it should be noted that this simulation assumes a flat ground plane. In rocky or irregular terrain, local structure from motion will add complex local variations to the underlying retinal motion patterns described here, and it remains to be explored how these complexities might influence locomotion. However, walkers have ample opportunity to learn the statistical patterns associated with locomotion over a variety of terrains, as well as the range of patterns associated with stable postures. It is likely that retinal motion cues from both eyes are combined to provide a robust estimate of both our movement through the 3D structure of the world, in what has been referred to as the “binoptic flow field” (Cormack, Czuba, Knöll, & Huk, 2017; Bonnen et al., 2020). Nonetheless, the monocular task-relevant features of retinal optic flow described here is robust to complex, sinusoidal and circular simulated eye trajectories (Videos 9), locomotion over flat terrain (Video 10), as well as the more circuitous routes taken in the rocky terrain (Videos 11, 12, and 13).

## 3 Discussion

We have demonstrated the ways that the optic flow patterns experienced during natural locomotion are shaped by the movement of the observer through their environments. The natural variation in heading during the gait cycle means that the flow pattern relative to the head is highly unstable. This is true whether the walker’s head (and eye) is directed towards a distant target or at the ground nearby to monitor foothold selection. The problem of heading perception is typically characterized as one of accounting for eye movements to recover the translational direction of the head. Some analyses posit an explicit representation of motion signals relative to the head (see Perrone, 2018; Royden et al, 1992, van den Berg and Beintema, 1997). In this case, the variation in the motion patterns relative to the head during the gait cycle make it unlikely that these signals could be used to control steering towards a goal. However, stabilization of gaze by the VOR means that retinal motion patterns during the gait cycle are much less variable and exhibit regularities that may be valuable for the control of locomotion. In particular, a walker can determine whether they will pass to the left or right of their fixation point by observing the sign and magnitude of the curl of the flow field at the fovea. In addition, the divergence map of the retinal flow field provides a cue for the walker’s over-ground velocity/momentum vector, which may be an essential part of the visual identification of footholds during locomotion over complex terrain. The magnitude of the image velocities relative to the head make it unlikely that instantaneous heading is useful for control of navigation involving steering towards a distant goal.

The geometric relationship between retinal optic flow patterns and the translation of the eye over a groundplane was carefully investigatedout in a series of papers by Koenderink and van Doorn (e.g. Koenderink, 1986; Koenderink & van Doorn, 1976, 1984). This work also suggested that retinal flow patterns might be informative, and our findings are consistent, with the large body of literature that shows how instantaneous heading can be computed directly from retinal flow (Longuet-Higgins & Prazdny, 1980; Heeger & Jepson, 1992; Lappe & Rauschecker, 1993; Perrone & Stone, 1994; Royden, 1997; Wang & Cutting, 1999; Perrone, 2018). A key focus of the present study is the analysis of how this pattern varies through the gait cycle. Much of the work on optic flow either implicitly or explicitly assumes that optic flow is primarily relevant to steering to a distant goal, and so the experimental simuli tend to use use constant velocity straight line motion that is poor represenation of the visual motion stimulus experienced during natural locomotion.Some work has suggested that heading estimates rely on information accumulated over time (W. H. Warren et al., 1991; Grigo & Lappe, 1999; Layton & Fajen, 2016). In partcular, recent work by Burlingam and Heeger (Burlingham & Heeger, 2020) demonstrates that instantaneous optic flow is insufficient for heading perception in the presence of rotation, and showed that heading judgments in the presence of rotation depend on the timevarying evolution of the flow (that is, the retinal acceleration field). Althrough, their stimuli did not mimic the kind of time variations generated by natural gait patterns, their finding that observers are sensitive to the temporal derivative of the retinal velocity field has an interesting relationship to the present discussion of divergence and curl. Divergence and curl are operators over V (pronounced “del”), which is defined as the 2 dimensional spatial gradient (derivative) of a vector field (in our case, the retinal velocity field). This means that our study examined ways to determine heading by way of the *spatial* derivative (∇) of retinal flow, where as Burlingham and Heeger solve a similar problem by way of the *temporal* derivitive (acceleration) of the same vector field. Thus, despite the obvious differences between the two approaches, both suggest that the solution to the determining heading from retinal flow lies in the consideration of the higher order structure of the instantaneous retinal flow field. Although it remains an empirical question as to whether and how either cue might be used in practice, redundancy is the key to robustness, so it is possible that both factors contribute to the visual control of natural locomotion.

In experiments that use a stationary observer and simulate direction on a computer monitor (Li & Cheng, 2013; Li & Niehorster, 2014), the strong sense of illusory motion accurate judgements of simulated heading indicate that humans are highly sensitive to full field optic flow. However, it does not necessarily mean that subjects use this information to control direction of the body when heading towards a distant goal. The complex, phasic pattern of acceleration shown here derives from the basic biomechanics of locomotion (Kuo, 2007). In the absence of direct measurements of flow during locomotion, the magnitude of the effect of gait has not been obvious. Thus it may have been implicitly assumed that the overall structure of optic flow during locomotion would be dominated by the effects of forward motion. This would require integration of head direction through the gait cycle, despite the large and rapid variation in direction, something that would require further investigation.

### 3.1 The use of optic flow during natural locomotion

Many of the primary challenges of human walking operate on much shorter timescales than steering towards a distant goal. Striding bipedalism is inherently unstable - successful locomotion requires a walker to constantly regulate muscle forces to support the mass of the body above the ground (McGowan, Neptune, Clark, & Kautz, 2010; Zajac, Neptune, & Kautz, 2002). In complex, rough terrain, these supporting muscle forces critically depends on the careful placement of the feet onto stable areas of terrain that are capable of supporting the walker’s body while redirecting the momentum of the center of mass towards future footholds (Matthis et al., 2017, 2018). Failure to place the foot in a stable location or failure to properly support the body within a step may result in injury. Given that walkers place their foot in a new location roughly twice per second, it is reasonable to assume that the underlying visual-motor control processes for foot placement and balance are fast, reliable, and efficient. It is also reasonable to suppose that subjects are able to learn the dynamic evolution of the flow patterns they generate during locomotion and so can use retinal flow patterns as a control signal. Indeed, it has been clearly demonstrated that optic flow exerts an important regulatory influence on the short-term components of gait. Expansion and contraction of optic flow patterns entrain the gait of subjects walking on treadmills, with the steps phase-locked to timing of expansion and contraction in the flow patterns (Bardy, Warren, & Kay, 1996, 1999; W. H. Warren, Kay, & Yilmaz, 1996). Additional research has shown that optic flow can speed up adaptation to an asymmetric split-belt treadmill (Finley, Statton, & Bastian, 2014). Other evidence indicates that the FoE had a strong influence on posture but not on heading(Schubert, Bohner, Berger, v. Sprundel, & Duysens, 2003). Salinas, Wilken, and Dingwell (2017) and Thompson and Franz (2017) have also demonstrated regulation of stepping by optic flow, and Reimann et al. (2018) has shown that optic flow perturbations produce a short-latency response in the ankle musculature. O’Connor and colleagues have shown that optic flow perturbations have a stronger effect when they are in a biomechanically unstable direction during standing and walking (O’Connor & Kuo, 2009), and that treadmill walkers will rapidly alter their walking speed to match altered optic flow before slowly adjusting back to their biomechanically preferred speed (O’Connor & Donelan, 2012). Taken together this body of research indicates a robust and sophisticated relationship between optic flow and the regulation of the short timescale biomechanical aspects of locomotion.

The data we present here do not directly demonstrate a role of retinal flow in gait control. However, all these studies suggest that the retinal optic flow cues described in this paper function as the control variables for stepping. In a study by Bossard et al (Bossard, Goulon, & Mestre, 2016), viewpoint oscillations consistent with head movements were added to a radial optic flow simulating forward self-motion. This was found to influence perception of distance travelled (as opposed to heading). This finding suggests that subjects are sensitive to the oscillatory effects of optic flow, and can integrate some aspects of flow information over the gait cycle and over longer time periods. However, this effect was only observed when the subjects were stationary, and not when they were walking on a treadmill (Bossard, Goulon, & Mestre, 2020). This suggests that the context - whether walking or not - is an important modulating factor in the way that flow is used.

### 3.2 Optic flow, visual direction, and the control of locomotor heading

The act of steering towards a goal does not necessarily involve use of optic flow. An alternative for control of heading in this sense is visual direction, Rushton, Harris, Lloyd, and Wann (1998) proposed that the perceived location of a target with respect to the body is used to guide locomotion, rendering optic flow unnecessary. Perhaps the strongest evidence for the role of optic flow in control of steering towards a goal is the demonstration by W. H. Warren, Kay, Zosh, Duchon, and Sahuc (2001) who pitted visual direction against the focus of expansion in a virtual environment, where walkers generate the flow patterns typical of natural locomotion. They found that although visual direction was used to control walking paths when environments lacked visual structure (and thereby lacked a salient optic flow signal), optic flow had an increasing effect on paths as environments became more structured. The authors interpreted this result to mean that walkers use a combination of visual direction and optic flow to steer to a goal when the visual environment contains sufficient visual structure. This is puzzling in the context of our findings, since the W. H. Warren et al. (2001) experiment used a fully ambulatory virtual environment, so the head-centered optic flow experienced those subjects would have had the same instabilities described here. How then can we reconcile these results?

In the W. H. Warren et al. (2001) experiment, the FoE and visual direction were put in conflict using a virtual prism manipulation that displaces the visual direction of the goal but not the FoE. However, in light of the present study, it seems that this prism manipulation would also induce a change in the retinal curl subjects experienced during forward locomotion. To see why, imagine a walker (without prisms) moving forward while fixating a point that is directly on their travel path. As described above, this walker would experience no net curl at their point of fixation. Now imagine the same situation for a walker wearing prism glasses that shift their visual environment 10 degrees to the right - In that case, an individual walking forward while fixating a point that is directly on their travel path would experience retinal flow consistent with fixating a point that was 10 degrees to their right. Therefore, the walker would experience a retinal flow field with counter-clockwise rotation (negative curl) as in Figure 5C and Video 6. If walkers use this retinal flow cue to control their foot placement, this effect might drive them to turn their body in the opposite direction within a step, in order to null this illusory retinal rotation/curl, resulting in a straighter path towards the goal. Put another way, retinal curl provides a moment-to-moment error signal that the walkers could use to actively correct for the effects of the prism. This hypothetical locomotor effect would be most pronounced in environments with salient optic flow, which could explain why the authors found that subjects walked in straighter lines in visually structured environments. In this interpretation, the walker’s act of directing gaze towards their goal provides an egocentric heading cue to help them steer towards their target while retinal optic flow provides an error signal that may be used to control their locomotion and balance on a shorter timescale. Thus the present focus on the retinal flow and its evolution over time may help unify research on longer-timescale control of steering towards a goal (Fajen, 2003) with the shorter-timescales associated with control of the mechanical aspects of balance and trajectory within a step.

### 3.3 Cortical involvement in the perception of optic flow

One of the insights from the observations in this study is that the stimulus input to the visual system is critically dependent on both the movement of the body and the gaze sampling strategies, especially in the case of motion stimuli. A similar point was made by Calow and Lappe (Calow & Lappe, 2008). Gaze patterns in turn depend on behavioral goals. In rough terrain gaze is mostly on footholds 2-3 steps ahead, whereas in regular terrain gaze is mostly directed to more distant locations (Matthis et al., 2018). This changes the pattern of flow on the retina as demonstrated. If humans learn the regularities in the flow patterns and use this information to modulate posture and control stepping, investigation of the neural analysis of optic flow ideally requires patterns of stimulation that faithfully reflect the patterns experienced during natural movement. A variety of regions in both monkey and human cortex respond to optic flow patterns and have been implicated in self motion perception, including MT, MST, VIP, CSv (Morrone et al., 2000; Wall & Smith, 2008; Yu, Hou, Spillmann, & Gu, 2017; X. Chen et al., 2013; Sunkara, DeAngelis, & Angelaki, 2015, 2016) and activity in these areas appears to be linked to heading judgments. However, there is no clear consensus on the role of these regions in the control of locomotion. For example, in humans, early visual regions and MT respond about equally to motion patterns that are inconsistent with self-motion (for example, when the FoE is not at the fovea) as they do to consistent patterns. MST had intermediate properties (Wall & Smith, 2008). Similarly in monkeys selectivity for ego-motion compatible stimuli was never total in a range of areas including MST and VIP (Cottereau et al., 2017). Strong, Silson, Gouws, Morland, and McKeefry (2017) also conclude that MST is not critical for perception of self-motion. Since a variety of different motion patterns have been used in these experiments, it may be necessary to simulate the consequences of self-motion with stimuli more closely matched to the natural patterns that take the kinematics of the body movements into account. Many neurons within MST respond vigorously to spiral patterns of motion, which is notable in light of the ubiquitous spiral patterns that appear in the present dataset (e.g. Video 3 and 12). Interestingly, although neurons in both MST and VIP signal heading, they do not strongly discriminate between the retinal patterns generated by simulated, as opposed to real, eye movements (Sunkara et al., 2015; Manning & Britten, 2019). This suggests that the information critical for heading is computed in a retinal reference frame (Sunkara et al., 2015; X. Chen, DeAngelis, & Angelaki, 2018). Similarly, Bremmer et al. (2017) used a decoding model to estimated heading from the MST of monkeys looking at flowfields and noted that their reliability dropped every time the monkeys made a saccade, indicative of a neural system meant to extract heading on a per-fixation basis. This result is consistent with the suggestion of the present paper, that the retinal patterns contain the critical information. The use of retinal flow for the control of movement requires that a walker has a notion of the body-relative location of their current fixation (Crawford et al., 2011), which might explain the well-established connection between vestibular cues and optic flow parsing (MacNeilage, Zhang, DeAngelis, & Angelaki, 2012).

## 4 Conclusion

Gibson’s critical insight on optic flow was that an agent’s movement through their environment imparts a structure to the available visual information, and that that structure can be exploited for the control of action (Gibson, 1950). In the intervening years, a large and fruitful body of research built upon this insight. However, the way optic flow is used by the visual system is difficult to intuit without direct measurement of the flow patterns that humans generate during natural behavior. Advances in imaging technology and computational image analysis, together with eye and body tracking in natural environments, have made it easier to measure and quantify these complex aspects of the visual input. Examination of the retinal flow patterns in the context of fixating and locomoting subjects suggests a change in emphasis and reinterpretation of the perceptual role of optic flow, emphasizing its role in balance and step control rather than in control of steering toward a goal. While many methods exist to compute instantaneous heading from the retinal flow field, a consideration of these patterns relative to the gaze point through the gait cycle provides a different context for the way the retinal flow information is used to control real-world, natural locomotion.

## 5 Acknowledgements

This work was supported by NIH 1T32-EY021462, R01-EY05729, and K99/R00-EY028229. Special thanks to John Cormack for his advice regarding the development of the streamline method for tracking the focus of expansion.

## 6 Methods and Materials

### 6.1 Experimental Subjects

Three human subjects participated in this experiment (1 female, 2 male; mean age: 28.7 +/−5 years, mean height: 1.79 +/− .14 m, mean weight: 78.3 +/− 18.8 kg, mean leg length: .96 +/− .68 m). One of the subjects was the first author of this manuscript. Subjects signed informed consent prior to participating in the experiment and all activities were approved by the Institutional Review Board at the University of Texas at Austin.

### 6.2 Equipment

Subjects’ gaze was tracked using a Pupil Labs mobile eye tracker (Pupil Labs, Berlin, Germany). Each eye was recorded at 120Hz with 640×480 resolution (the eye cameras), while an outward facing camera mounted 3cm above the right eye (the world camera) recorded at 30Hz with 1920×1080 resolution and a 100degree diagonal field of view. Subjects’ eyes were shaded using roll-up optometrist sunshades that covered the eyes and eye cameras but left the world camera uncovered. This method of shading the infrared eye cameras from sunlight was less robust than the full IR blocking face shield used in (Matthis et al., 2018), but was necessary to prevent the computational video analysis algorithms (described below) from being affected by reflections on the inside of the mask. For this reason, data collection was conducted during a time of day when the walking path was mostly shaded from direct sunlight. Kinematics were recorded using the Motion Shadow full body motion capture system using inertial measurement units recording at 100Hz. Raw data were initially recorded on a backpack-mounted laptop worn by the subject and later post-processed using custom code written for Matlab (MathWorks, Natick, MA, USA).

### 6.3 Data and code availability

In the fullness of time, all raw data will be posted in an online repository along with the Matlab code necessary to recreate each of the figures and videos in this manuscript.

### 6.4 Experimental Task

Subjects walked along two separate paths in the greenbelt of Austin TX, USA-A flat, tree-lined path consisting of mulched woodchips in the Shoal Creek trail in Pease District Park (the “woodchips path”) and a rough, rocky creekbed consisting mostly of large boulders (the “Rocky path”). The woodchips path was selected because it was flat enough that foot placement did not require visual guidance, but was visually textured enough for the optical flow detection algorithms to detect visual motion (in contrast to something like a concrete sidewalk). The rocky path was the same path used in the Rough Terrain condition of Matthis et al. (2018).

Subjects walked from start to the end of the Woodchips path while following one of three sets of instructions - Free walking, Ground looking, and Distant Fixation. In the Free Walking condition, subjects were instructed to simply walk from the start to the end location without any explicit instructions for what to do with their eyes. In the Ground Looking condition, subjects were asked to walk while looking at the ground at an approximately fixed distance ahead. In the Distant Fixation condition, subjects were asked to walk while maintaining fixation on a self-selected distant object that was roughly at eye height (switching targets as necessary to maintain distant fixation). This condition was intended to most closely match the psychophysical tasks involved in previous research on optic flow during locomotion, i.e. fixation on a distant object while moving forward without head movement. Locomotion in the Rocky trail was challenging enough that subjects were not asked to complete a secondary task. They were simply asked to walk from the start to end position at a comfortable walking pace.

Subjects completed three out-and-back walks in the woodchips path, for a total of 6 trial/walks in that condition. There were two repetitions of each condition in the woodchips (one per walking direction). Subjects completed 4 out-and-back walks on the Rocky path, for total of 8 trial/walks. Because the woodchips path was significantly longer than the rocky path, a similar amount of data was collected in each condition.

### 6.5 VOR-based calibration and data postprocessing

We used a procedure analogous to the VOR-based calibration method developed in Matthis et al. (2018), with some alternations due to the change in eye tracker. The Pupil Labs tracker used in this study estimate gaze for each eye using 3D spherical eye models generated within the coordinate frame of each eye camera. Using the procedure described below, the gaze estimates for each eye were rotated to align with the reference frame of the full-body kinematic estimates from the IMU-based motion capture system.

The calibration procedure was completed at the start of data recording for each terrain condition for later processing. Subjects stood on a calibration mat that had marks for the subjects’ feet, a high visibility marker located 1.4 m ahead of the vertical projection of the midpoint of the ankle joint, and a 0.5 m piece of tape at the same distance. Following the experimenter’s instruction, subjects maintained fixation on that point while slowly moving their heads up/down, left/right, and along the diagonals. In addition to help determine the subject’s 3D gaze vector by relating eye and head movements, data from this portion of the record were used to calibrate the eye tracker (similar to the “head tick” method described in Evans, Jacobs, Tarduno, and Pelz (2012), except that our subjects moved their heads smoothly).

The post-processing procedure to determine the subjects’ world-centered 3D gaze vector is described below. Note that because each eye was calibrated independently, so any alignment between the gaze vectors of the right and left eye is an indication that the calibration was completed accurately.

1. Align timestamps of Pupil eye tracker and Shadow IMU data. Because the eye tracker and the motion capture system were being recorded on the same backpack mounted laptop, the timestamps from the two systems were already synchronized. Aligning the two data streams was a relatively simple matter of ensuring that both systems calculated time relative to some external temporal reference (rather than some unknown internal reference, such as “computer boot time”).
2. Resample data from eye tracker and IMU to ensure constant 120Hz framerate. Kinematic data from the Shadow system (recorded at 100Hz) were upsampled to 120Hz to match the framerate of the eye camer as. Foll owing this, the left eye, right eye, and kinematic data streams were resampled to synchronize the three data streams.
3. Estimate eye center coordinates relative to head position estimates from Shadow system. This location will serve as the center for the spherical eye model generated by the Pupil Labs eye tracker.
4. Situate 3D spherical eye model from Pupil tracker onto eye center estimate calculated in step 3. Because this eye model was generated in the reference frame of the relevant eye camera, gaze vector orientations will be arbitrary when placed in the world-centered reference frame of the kinematic data from the Shadow system. The next step will align these gaze vectors with the subejcts’ world-centered gaze.
5. Use data from VOR calibration task to rotate gaze data from Pupil tracker to align with the subjects’ world-centered gaze using unconstrained optimization (MATLAB’s fminunc function).
  a. Begin with a starting guess where the Euler angle rotation of the gaze data is [0 0 0].
  b. Rotate all gaze data by this rotation, and then rotate each gaze vector by the subject’s head orientation on the frame that it was recorded.
  c. Calculate intersections between each gaze vector and the ground plane. If a gaze vector does not intersect the ground plane, truncate it at 10 m.
  d. Calculate error of this camera alignment rotation, defined as the mean distance between the intersection point of each gaze vector and the calibration point that subjects were fixating (located 1.4 m ahead of the vertical projection of the subject’s ankle joints).
  e. Use fminunc to minimize this error by optimizing the Euler angle rotation to apply to the gaze vectors prior to applying the rotation specified by the subject’s head orientation. The correct orientation will cause subjects’ head rotations to cancel their eye movements to maintain the gaze vector alignment with the calibration point (that is, the correct gaze alignment rotation will preserve VOR-based eye compensation for head rotation).
6. Once the correct gaze alignment rotation has been identified, apply it to all subsequent gaze data from this recording prior to rotating each gaze point according to the subject’s head orientation on that data frame.

### 6.6 Video Analysis and optic flow estimation

We used the Matlab Camera Calibration toolbox to estimate the lens intrinsics of the head-mounted world camera of the Pupil tracker (3 radial distortion coefficients with skew correction). This method utilizes a checkerboard of known size to determine the distortion caused by the wide angle lens of the camera. This calibration effectively allows us to treat the images from the head mounted camera as if they were recorded by a linear pinhole camera. We then estimated the visual motion on each recorded frame of the head-mounted camera using the Deep Flow optical flow estimation algorithm(Weinzaepfel et al., 2013) in OpenCV (Bradski & Kaehler, 2011). DeepFlow is a dense optical flow estimation algorithm that provides a motion estimate at each pixel of each frame of the analyzed video. Despite its name, this method does not rely on deep learning methods to detect optic flow. Internal testing found that deep-learning based methods are prone to biases (likely due to their training data), and so are not appropriate for use as a scientific research method.

#### 6.6.1 Estimating retinal reference frame

In order to estimate the visual stimulus in a retinal reference frame, each recorded frame from the undistorted world camera was projected onto a sphere and placed so that that subjects’ point of regard on that frame was aligned with 0,0 in the retinal reference frame. To account for inaccuracy in this eye tracker, fixations were “idealized” by adjusting the placement of the image to null any residual motion detected at that point of fixation. (Inspection of the videos revealed only minimal residual motion during fixation.) In this way, the retinal reference frame videos provide an estimate of the visual motion that was incident to the subjects’ retina, assuming “perfect” fixation and a spherical pinhole camera model of the eye.

### 6.7 Optic Flow simulation with a spherical pinhole camera model of the eye

We created a geometric simulation to provide a more nuanced picture of the way that the movement of the body shapes the visual motion experienced during natural locomotion. To estimate the flow experienced during various types of movements, a simulated eye model was generated using the following procedure. Most of the geometric calculations used in this model rely heavily on the Geom3D toolbox on Mathworks.com (Legland, 2020)

1. Define the groundplane as an infinite, flat plane with zero height.
2. Define an evenly spaced grid of points on the ground plane, which will eventually be projected onto the back of the simulated retina.
3. Define the spherical eyeball as a sphere with a radius of 12 mm (the anatomical average radius of the human eye). In this model, one pole of the sphere is defined as the ‘pupil’ and the opposite pole is defined as the ‘fovea’. Place the center of this eye model at the correct location in 3D space. This location is either determined from the recorded motion capture data (as in Videos 10 and 13) or by determining a prescribed path for the eye to follow (as in Videos 4 - 9)
4. Rotate the eye model to face the fixation point on that frame. Orient the eye so that there is a line passing through so that a line passing through fovea and pupil will also pass through the fixation point on the ground. This rotation was defined so as to minimize torsion about the optical axis.
5. Define the field of view of the eye by projecting a cone from the pupil determining the intersection points between this cone and the flat ground plane. For this study, we defined the field of view cone to have a 60 degree radius..
6. Project all the groundpoints within the field of view through the pupil and onto the back of the 3D spherical retina.
7. Resituate projected points onto a 2D polar plot where the vertical axis (pi/2) defines the anatomical superior of the eye, and the eccentricity is defined as the “great circle” distance of the projected point from the ‘fovea’ of the spherical eye. Keep track of these projected locations across successive frames, to allow for calculation of optic flow on later steps.
8. Calculate flow per frame by determining the distance between the projection of each point on successive frames. The flow on the first frame is defined to be zero everywhere. If a particular point was not within the eye’s field of view on the previous frame, flow at that point is undefined on the first frame that point became visible.
9. Use the scatteredInterpolant function in Matlab to define an evenly spaced vector field across the retina that shares the structure of the projected-dot flow field determined in the previous step. This step is necessary to be able to calculate the Curl and Divergence of the vector field.
10. Calculate divergence and curl of this evenly spaced retinal velocity grid using the divergence and curl functions in Matlab.
11. Spend two years fastidiously creating overcomplicated videos to highlight various features of the resulting flowfields (optional)

#### 6.7.1 Estimating the Focus of Expansion

To estimate the location of the focus of expansion on each frame, each frame from world camera was first processed by the DeepFlow optical flow algorithm described above. This method provides a motion estimate for each pixel of the video frame, providing a 2 dimensional vector field with the same dimension as the original video for each recorded frame. To track the Focus of Expansion (FoE) in each frame, this vector field was first negated (all vectors were multiplied by −1), which effectively transforms the FoE from a repellor node (vectors pointing away from the FoE) into an attractor node (vectors pointing towards the FoE). Then, a grid of particles was set to drift on this negated flow field using the streamlines2 function in Matlab. The paths traced by these particles provide information about the underlying structure of the optic flow on each frame, represented as white lines in Figures and Video. These streamlines represent the line integrals of the optic flow vector field measured on each frame.

The location of the FoE was determined by keeping track of the final fate of each of the drifting particles and tagging the location where the majority ended up as the pixel location of the FoE on that frame. On frames where fewer than 50% of particles could be found in a particular location on the video screen, it was determined that the FoE was not in the field of view of the camera on that frame (this could be verified visually by noting that the streamlines on those frames do not converge to a single point). To calculate the velocity of the FoE on each frame, we calculated the 2D distance traveled by the FoE on successive frames, multiplied by that by the field of the view of the camera (to convert to Degrees Visual Angle) and then divided by the framerate (to convert to Degrees per second).

To ensure that this method was capable of detecting a stable FoE (that is, to ensure that the high velocity of the FoE that we record was not just an artifact of the detection method), we applied this analysis to the video from a DJI Phantom 4 quadcopter as it traveled along a straight horizontal and vertical flight path. The camera of this quadcopter is stabilized by a 3-axis gimbal mount, which is essentially a mechanical equivalent of the VOR-reflex. The resulting FoE is nicely stable, indicating that this method is capable of detecting a stable, low-velocity FoE in a video, if it is present (Video 14).

## Footnotes

1 Note that although previous research has suggested that walkers may engage in “travel gaze” during locomotion (where gaze is “visually anchored in the front of the individual and carried along by the whole body movement (Patla & Vickers, 2003)“), this behavior is inconsistent with gaze stabilization reflexes such as the VOR and OKN. Consequently, the Ground Looking condition actually consisted of a series of brief fixations and small saccades to keep gaze roughly a fixed distance ahead of the walker.

2 If gaze is not perfectly stationary during fixation, there will be a small translational component added to the retinal motion. Unfortunatley, a full explication of this effect is beyond the scope of this paper, as the error associated with incomplete gaze stabilization during fixation is of a similar magnitude to the errors expected from the current generation of mobile eye trackers. However, we estimated the magnitude of this slip and found that stabilization during a fixation is quite good, with the median slip being less than a degree of visual angle along the direction of travel during a fixation, with the gain being slightly less than 1.0.For this reason, we chose to ‘‘idealize” fixations by nulling motion at the fovea and kept our research questions to a scale that will not be sensitive to this simplification.

